# Collective polymerase dynamics emerge from DNA supercoiling during transcription

**DOI:** 10.1101/2021.11.24.469850

**Authors:** Stuart A. Sevier, Sahand Hormoz

**Affiliations:** Department of Systems Biology, Harvard Medical School, Boston, MA 02115, USA; Department of Data Sciences, Dana-Farber Cancer Institute, Boston, MA 02215, USA; Broad Institute of MIT and Harvard, Cambridge, MA 02142, USA

## Abstract

All biological processes ultimately come from physical interactions. The mechanical properties of DNA play a critical role in transcription. RNA polymerase can over or under twist DNA (referred to as DNA supercoiling) when it moves along a gene resulting in mechanical stresses in DNA that impact its own motion and that of other polymerases. For example, when enough supercoiling accumulates, an isolated polymerase halts and transcription stops. DNA supercoiling can also mediate non-local interactions between polymerases that shape gene expression fluctuations. Here, we construct a comprehensive model of transcription that captures how RNA polymerase motion changes the degree of DNA supercoiling which in turn feeds back into the rate at which polymerases are recruited and move along the DNA. Surprisingly, our model predicts that a group of three or more polymerases move together at a constant velocity and sustain their motion (forming what we call a polymeton) whereas one or two polymerases would have halted. We further show that accounting for the impact of DNA supercoiling on both RNA polymerase recruitment and velocity recapitulates empirical observations of gene expression fluctuations. Finally, we propose a mechanical toggle switch whereby interactions between genes are mediated by DNA twisting as opposed to proteins. Understanding the mechanical regulation of gene expression provides new insights into how endogenous genes can interact and informs the design of new forms of engineered interactions.

PACS numbers:

---

Numerous physical processes contribute to gene expression [1]. For example, transcription, an essential step in gene expression where DNA is converted to RNA has a mechanical component. During transcription, an RNA polymerase can twist DNA and generate stresses on DNA [2, 3] that impact its own motion and that of other polymerases [4] affecting gene expression [5]. Therefore, to understand the dynamics of gene expression, we need to account for the mechanical nature of transcription [6].

Gene expression occurs in a stochastic (‘bursty’) manner [7]. Early insights into gene expression fluctuations [8–12] led to phenomenological frameworks that did not account for DNA mechanics or the physical forces involved in transcription [13]. These models were followed by theoretical [14–16] and experimental studies [17–19] that found over and under twisting of DNA, or DNA supercoiling, to be a robust mechanism of generating transcriptional bursting. In addition to influencing fluctuations, experimental [20] and theoretical [14] observations have shown that DNA twisting can halt isolated polymerases and stop transcription whereas multiple polymerases can undergo effective elongation [4] and influence the recruitment of polymerases to neighboring promoters [21]. Despite the advances that these studies have made, we still lack a comprehensive framework that captures how DNA twisting by RNA polymerase during transcription feeds back into the recruitment and motion of other RNA polymerases which in turn change the degree of twisting of DNA.

Here, we construct a model of mechanical aspects of gene expression (referred to as mechanical epigenetics) that captures the feedback cycle between DNA twisting and RNA polymerase recruitment and motion. Within our framework, multiple polymerases interact non-locally via twisting of DNA generated by the movement of the polymerases along the gene. We show that these interactions play a key role in setting the velocity at which polymerases can move along a gene which in turn sets the degree of gene expression fluctuations. Surprisingly, we demonstrate that three or more interacting polymerases undergo sustained motion whereas isolated polymerases are halted by mechanical forces (a collective phenomenon that we call a polymeton). We also show that incorporating the impact of DNA twisting on both recruitment of RNA polymerase and their interaction can correctly predict observed fluctuations in gene expression. Finally, we use the mechanical coupling between polymerases to propose a computational model of a toggle switch whereby interactions between two genes are mediated by DNA twisting as opposed to proteins.

## Model description

Two major factors determine gene expression output. The first is the rate at which polymerases are recruited at the transcription start site (TSS), referred to as the initiation rate. Second is the rate at which the recruited polymerases are transported from the start site to the termination site, referred to as the elongation rate. The rates of initiation and elongation together determine the rate of gene expression output.

Fluctuations in the output are not necessarily equal to fluctuations created during initiation because polymerase velocity and spacing can change during transport. To understand the fluctuations in the output and how it relates to the fluctuations in the input, we need to model the transport of polymerases and quantify their contribution to the fluctuations in the output.

Our model describes the position and velocity of the polymerases during transport. We assume that there are *N* polymerases between the transcription start site and the termination site. The ith polymerase has position *x*_*i*_ and velocity *v*_*i*_. The density function of the polymerases *ρ* = Σ_*i*_*δ*(*s* − *x* (*t* _*i*_)) encodes the position of all the polymerases along the gene, where position along the gene is parameterized by *s* (in bp). Similarly, the flux of polymerases along the gene is defined as:

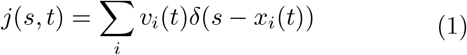

Flux *j* at position *s* corresponds to the number of polymerases crossing *s* per unit time. The unit of flux is the same as that of the initiation and output rates. The flux captures the transport of polymerases along the gene and can be used to link initiation and elongation to output (Figure 1). At steady state, the initiation rate is equal to the output rate when averaged over a sufficiently long period of time. In addition, these rates should equal the average flux at any point along the gene when there is no depletion of polymerases along the gene. However, the fluctuations in these three rates are not necessarily equal. To understand how variations in the flux control output fluctuations, we need to incorporate the physical factors that modify polymerase velocity and the initiation rate.

**FIG. 1:**
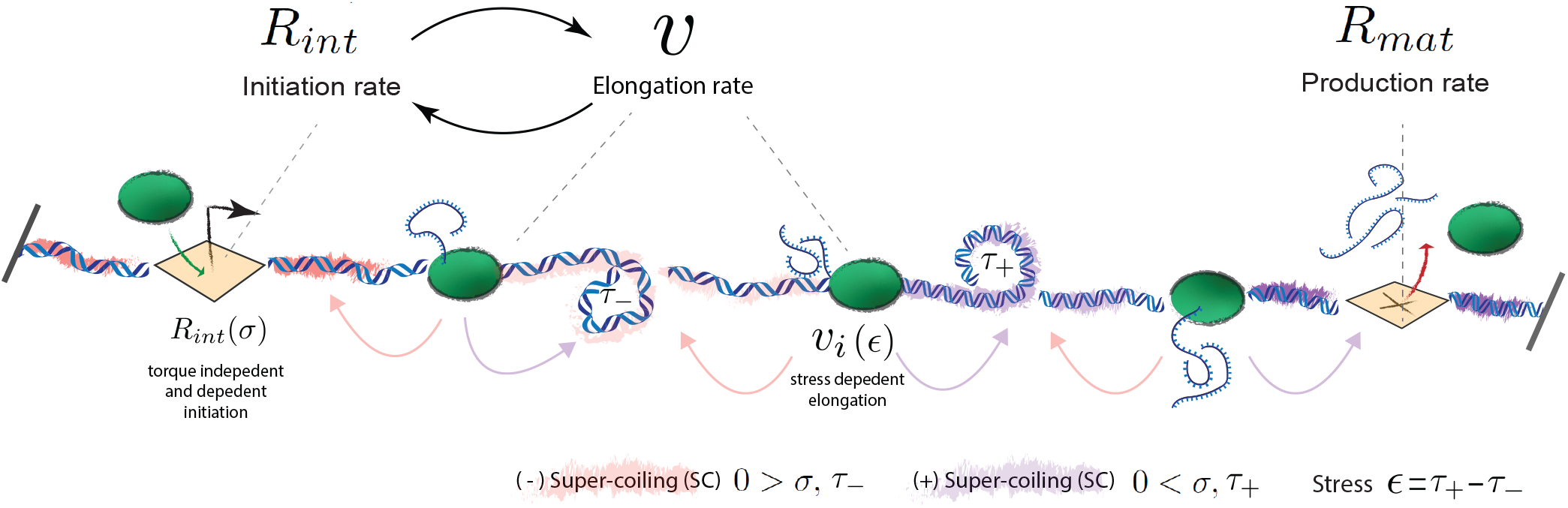
Schematic of the model of the role of supercoiling during transcription. Gene expression output is determined by both polymerase recruitment (initiation) as well as polymerase transport along the gene (elongation). During elongation, polymerases produce supercoiling, the over and under twisting of DNA (shown in purple and red respectively), causing a corresponding change in the torque and in turn the torsional stress on the DNA. DNA supercoiling and torque can be transported between polymerases creating non-local interactions between polymerases that influence elongation and initiation. These interactions also impact both the average rate of mRNA production and its fluctuations.

DNA supercoiling (SC), the over and under-twisting of DNA [22], can control both the initiation rate and the velocity [4, 5] of polymerases. Importantly, as polymerase moves along the gene it can change the degree of supercoiling. This is because to transcribe, a polymerase has to either (1) rotate to follow the helical grooves of DNA and/or (2) DNA itself has to twist as it’s pulled through a polymerase [22]. Polymerase rotation does not change the degree of supercoiling whereas DNA twisting does [14]. Therefore, we need to incorporate the feedback between polymerase velocity and DNA supercoiling in our model. Following a physical construction of the twin-domain model of transcription [14], we define 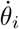 to be the rate of rotation (angular velocity) of the ith polymerase. *ϕ*(*s*) is the degree of twisting of DNA at position *s*. We define *ϕ*(*s* = 0) = 0. Even with zero supercoiling, as *s* increases *ϕ* also increases because of the natural helical form of DNA at a rate of *ω*_*o*_ = 1.67*rad/bp* (for relaxed DNA). 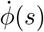 is the rate of twisting per unit time at position *s. ∂*_*s*_*ϕ* evaluated position *s*_0_ captures how the degree of twist changes when moving along the gene from position *s*_0_ to a point infinitesimally away *s*_0_ + *ds. ∂*_*s*_*ϕ* is referred to as the local twist density.

Thus, *v∂*_*s*_*ϕ* determines the rate at which a polymerase encounters twist when moving at velocity *v* along the gene. To follow the grooves of DNA, a combination of polymerase rotation or DNA twisting (in the opposite direction) must occur [14]. Therefore, for the ith polymerase, the equation

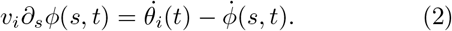

encodes the local interplay between DNA twist density encountered during elongation, polymerase rotation and further DNA twisting.

The supercoiling density *σ* is defined as the change along the gene of the over or under twisting of DNA past its natural twist density (*ω*_0_ = 1.67(*rad/bp*)) which can be expressed as

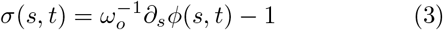

We can write equation 2 using the supercoiling density *σ* instead of the local twist density *∂*_*s*_*ϕ*.

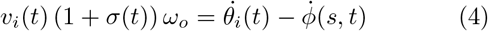

We can use mechanics to relate the rate of rotation of polymerase 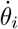 and the rate of twisting of DNA 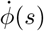 to the torque applied to the polymerase or DNA at position *s*. At each position *s* there exists some amount of torque *τ* (*s*) in response to the amount of twisting *ϕ*(*s*) at that position. The torsional stress *ϵ*_*i*_(*t*) = *∂*_*s*_*τ* (*s* = *x*_*i*_, *t*) (also called the local torque per unit length) exerts a damping force on the polymerase that rotates it. Intuitively torsional stress can be thought of as the difference of the torque in front and behind the polymerase. Specifically, the rate of rotation of ith polymerase is given by,

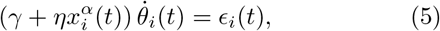

where the drag coefficient has a constant term *γ* and a term that increases with the distance *x*_*i*_ of the polymerase from the transcription start site. The drag coefficient increases with the distance of the polymerase from the start site because the nascent RNA attached to the polymerase increases in length as the polymerase moves along the gene. *γ* captures the drag of polymerase and DNA without nascent RNA attached. *α* is a phenomenological parameter that captures how the drag coefficient changes with the distance from the start site [23].

Similarly, the rate of rotation of DNA at position *s* away from transcription is related to the torsional stress experienced by the DNA at that position.

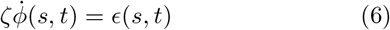

where *ζ* is the constant drag coefficient of DNA and *ϵ*(*s, t*) = *∂*_*s*_*τ* (*s, t*).

By substituting equation 5 into equation 4 to eliminate 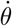 we obtain an equation for the rate of rotation of DNA at the position of the ith polymerase

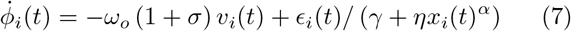

This equation directly relates the dynamics of DNA twisting to the rate of polymerase elongation *v*_*i*_ and positioning *x*_*i*_.

To write equation 7 for any position *s* on the gene, we combine the discrete drag coefficients of individual polymerases and the continuous drag coefficient of DNA and define,

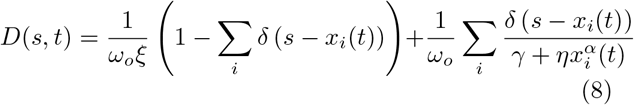

Using the above form of the the drag coefficient and our previous definition of flux j (Eq.1), we can write down an equation for the dynamics of DNA twisting that combines the contribution of polymerases with that of DNA itself

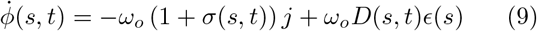

By applying equation 3 we arrive at a supercoiling density transport equation for transcription

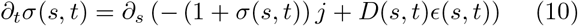

This result stands in contrast to previous models [16, 24] which assume a constant diffusion coefficient and neglect the fact that polymerases serve as both a source of supercoiling as well as barriers to its free diffusion [25].

We can write equation 10 explicitly in terms of *σ* (and thus stress) by specifying the local torque as

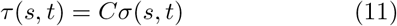

where *C* is the twist modulus of DNA (75*nm · k*_*b*_*t* [26]). Here we do not account for DNA bending or writhing in our simple model though their analytical inclusion is straightforward [27–30]. Their inclusion can lead to DNA buckling resulting in altered twist transport and torsional responses. To account for buckling in our simulations, we will utilize a phenomenological relationship between torque and supercoiling (see S.I.).

The flux term in the above equation and the diffusion constant depend on the position and the velocity of the polymerases. To close the set of equations in our description, we need to relate the position of the polymerases to their velocity and relate their velocity to the supercoiling density *σ*. The position of the polymerase ith change in time as 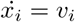.

We assume a phenomenological relationship between the supercoiling density and the velocity of polymerase *i* using the following functional form

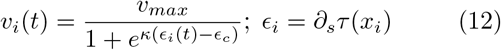

where *ϵ*_*i*_ is the torsional stress experienced by the *ith* polymerase as previously defined. *ϵ*_*c*_ the torsional stress cut-off (around 0.2*N* for E.Coli [20] assuming that the length scale of the polymerase is 0.6 angstroms [31]). *v*_*max*_ is the maximum velocity that polymerases can travel when *ϵ*_*i*_ << *ϵ*_*c*_. Conversely, if *ϵ*_*i*_ >> *ϵ*_*c*_ then the polymerase stalls. This phenomenological form is motivated by the empirical observations made in [20].

Taken together, equations eqs. (10) to (12) provide a closed-form description of the generation and transport of supercoiling density along the gene due transcription as well as the position and velocity of each polymerase. To solve these equations, we need to also specify the initial conditions and the boundary conditions. *s* = 0 and *s* = *L* are the positions of the two boundaries of the system. The boundaries of the gene itself (marked by the position of the transcription start site and termination site) are contained within the boundaries of the system. For an open system (DNA that is free to rotate at the boundaries) we have *σ*(*s* = 0) = *σ*(*s* = *L*) = 0. Similarly, for DNA that forms a closed loop *σ*(0) = *σ*(*L*), *∂*_*s*_*σ*(0) = *∂*_*s*_*σ*(*L*) so that the supercoiling density and torques match at the beginning and end of the system but are not necessarily zero. In our simulations, we use fixed boundary conditions where the gradient of supercoiling density at the two boundaries is zero *∂*_*s*_*σ*(*s* = 0) = *∂*_*s*_*σ*(*s* = *L*) = 0 at all times but the supercoiling densities are not necessarily equal to each other *σ*(0) ≠ *σ*(*L*).

The above equations with the boundary conditions specified can be simulated to study the generation and transport of supercoiling density along the gene. In our simulations, we assume the limit of fast diffusion of supercoiling density between the polymerases. This limit corresponds to *ζ* → 0 in Equation 8. With this assumption, the supercoiling density is constant along DNA regions between polymerases allowing for us to directly use equations 7 to keep track of supercoiling in alignment with previous efforts [15].

Finally, we incorporate the role of topoisomerases into our simple model. Topoisomerases play an important role in regulating transcription in both bacteria and eukaryotes [32]. General classes of topoisomerases are formed by their ability to relieve either positive or negative supercoiling density as well as their mechanisms of actions which use either single or double-strand breaks to modify the supercoiling density [33].

If no mechanism to relieve supercoiling is included (such as the one provided by topoisomerase) large amounts of supercoiling density accumulate and stalling occurs for reasonable choices of parameters as observed in experiments [34]. To incorporate topoisomerase action in our model, at random time-points, we scale the supercoiling at every point along the gene by the same constant factor by re-assigning the twist at each polymerase *ϕ*_*i*_ → *aϕ*_*i*_ (*a* = 0.1 in our simulations. The value of this constant is not important because the rate of topoisomerase action is the free parameter in our simulations).

The rate at which topoisomerase acts is set by the difference of the supercoiling density at the the two boundaries:

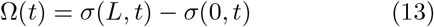

Ω(*t*) roughly corresponds to the accumulation of supercoiling density along the gene over time. In our simulations, we use the following form for the rate of topoisomerase action as a function of Ω(*t*):

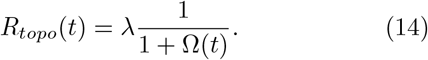

Thus, the overall rate of supercoiling density removal due to topoisomerase action goes as

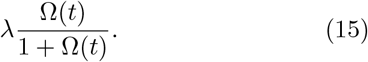

so that the rate of removal saturates with increasing levels of supercoiling at a fixed topoisomerase concentration. The above form is motivated by empirical observations of the dynamics of topoisomerase action in vitro [35]. In these experiments, supercoiled plasmids were prepared at varying concentrations effectively altering the amount of supercoiling present Ω [35]. Multiple types of topoisomerases were then added to remove the supercoiling. All topoisomerases displayed kinetic behavior in the form of equation 15. Thus, the phenomenological action of topoisomerase in our model closely captures the known kinetic properties of topoisomerases. As a check, we tried multiple alternative phenomenological forms relating Ω(*t*) to the rate of topoisomerase action: a constant rate independent of Ω(*t*); a rate proportional to Ω(*t*); and a rate that approaches zero with increasing Ω(*t*). The form used in equation 15 was the only one that was consistent with empirical observations of how topoisomerases regulate polymerase velocities and fluctuations in mRNA production (see S.I.).

### Simulations

We simulated the model described above to understand how initiation, transport, and supercoiling work together to determine gene expression output (the simulations are described in detail in supplemental section A). To do this, we simulated a single isolated gene of length 1000 bp contained within a total stretch of DNA of length *L* = 3000*bp*. The start site of the gene is located at *s* = 1000*bp*. At *s* = 0 and *s* = *L* DNA will be prevented from freely rotating causing supercoiling density to build up at the boundaries corresponding to the fixed boundary condition.

We begin with a constant initiation rate that does not depend on the supercoiling density at the transcription start site. Therefore, the loading process of polymerases at the transcription start site (*s* = 1000*bp*) is a Poisson process with rate *R*_*int*_. We will later consider an initiation rate that is a function of the supercoiling density at the transcription start site.

As the polymerases move along the gene, the supercoiling density changes according to equation 7 which governs the local twist change at each transcription site. We set the drag coefficient *γ* = 10^−1^[*pNs*], and the phenomenological parameters *η* = 10^−4^[*pNs/bp*^2^] and *α* = 2. While there is little empirical data to determine the precise values of these parameters, biophysical considerations (as well as the observation that short genes do not induce supercoiling while long ones do [25]) implies a drag greater than 1*pNnm* for a nascent transcript of length 1*kbp* or greater rotating at 10*rad/s*. We will regardless show that our results are robust to changes in the values of these parameters (see Discussion and S.I.). We assume that the torque is related to the supercoiling density using the functional form shown in S.I. section C. This choice is motivated by empirical observations [26] and only contains one free parameter which sets the torque at which DNA buckles. We also ensured that our results are robust to the choice of this parameter (see Figure S1).

When a polymerase reaches the transcription termination site, a mature mRNA is produced that then is removed at a constant rate *μ* = 10^−2^*s*^−1^. The simulations were started with no polymerases on the gene and ran for a sufficiently long period of time to reach steady-state when the number of RNA polymerases stabilized. We observed that the simulations reached steady-state typically after 10 minutes out of a total of 1 hour of simulation time (time-scale set by *μ*).

Figure 2a shows a snapshot of the simulation with 3 polymerases moving along the gene. For each simulation run we computed the number of mRNA molecules averaged across multiple simulation samples after each simulation reached steady state. The average number of mRNA is plotted as a function of the initiation *R*_*int*_ for different values of parameters *λ* and *v*_*max*_ (Figure 2b). As expected the number of mRNA molecules is proportional to *R*_*int*_ but does not depend on the values of *λ* and *v*_*max*_. This is because at steady state the rate at which polymerases are loaded must equal to the rate of production of mRNA.

**FIG. 2:**
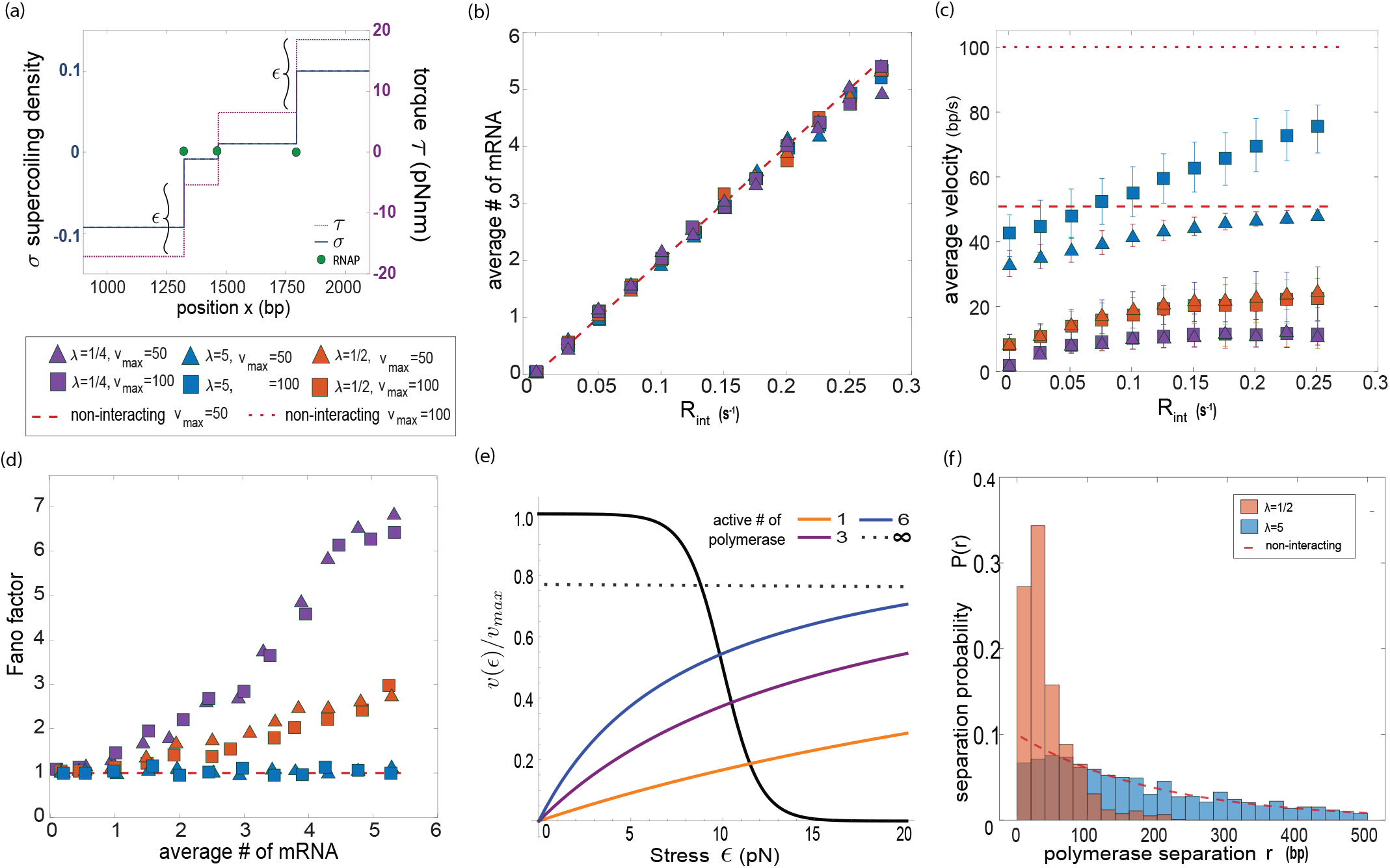
DNA supercoiling mediated interactions between polymerases do not alter the average mRNA production rate but link polymerase clustering to mRNA bursting.(a)Transcribing polymerases generate supercoiling density *σ* and accompanying torque during elongation. (b)The average number of mRNAs produced is insensitive to the elongation kinetics of the polymerases and is solely determined by the initiation rate. Changing the elongation kinetics by altering the maximum polymerase elongation rate *v*_*max*_ or the rate of topoisomerase action *λ* does not change the production rate.(c)Average elongation rates depend on the initiation rates and therefore the average number of polymerases present on the gene, demonstrating a cooperative interaction between the polymerases. The elongation rate plateaus to a value that is predominantly set by the rate of topoisomerase action. (d) A monotonic relationship between the mean mRNA production rate and its fluctuations (Fano factor) emerges with the slope determined by the rate of topoisomerase action. High rate of topoisomerase action results in fluctuations that resemble the Poisson statistics of non-interacting polymerases (red dashed line). (e) Analytical expressions for the elongation rate and the rate of topoisomerase action as a function of the stress. The rate at which supercoiling is added is proportional to the elongation rate. The steady-state value of stress and in turn the elongation rate is set when the rate of addition of supercoiling equals the rate of its removal by topoisomerase 17 (f) Supercoiling mediated interactions change inter-polymerase separation distances. The separation distance between the polymerase nearest the TSS and its closest neighbor, *r*, decreases with increasing accumulation rate of supercoiling density. The altered separation distances result in higher fluctuations in gene expression. The distribution of separation distances deviates from that of non-interacting polymerases that follow Poisson statistics (exponentially distributed separation distances, shown by red line). Simulation details described in Supplemental Section A.

The average velocity at which the polymerases move along the gene (shown in Figure 2c) also depends on *R*_*int*_ but saturates to a value that does not necessarily correspond to *v*_*max*_, especially when the rate of topoisomerase action *λ* is low or the ends of DNA are free. Importantly, this behavior recapitulates three empirical observations. One is that the polymerase velocity changes as the rate of topoisomerase action changes [4, 17]. Second is that the velocity of polymerases can be smaller than the bare velocity (*v*_*max*_/2) defined as the velocity of a single polymerase moving along a linear piece of DNA with open boundaries where there is no accumulation of supercoiling density [4, 17]. Third, actively elongating polymerases can be slowed by turning off further polymerase initiation [4] (Figure S3).

To gain an intuition for this behavior, consider the simplest case where all the polymerase velocities are equal to *v*. In addition, we assume that there is sufficiently large drag (*γ* + *ηx*^*α*^ >> 1) on each polymerase so that we can ignore polymerase rotation 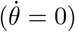. In this case, the supercoiling density generated by each polymerase is canceled by the supercoiling density generated by the neighboring polymerases except for at the boundaries where there is no cancellation. Then the rate of supercoiling density generation is −*ω*_*o*_*v* at the boundary closest to the start site and *ω*_*o*_*v* at the other boundary (Equation 10). Therefore, we can write down an equation for the rate of change Ω (eq.13) as

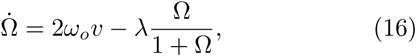

where the second term on the right-hand side is the rate of removal of supercoiling density by topoisomerase action. At steady state,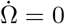.

If we make the additional simplifying assumption that the change in supercoiling density across each polymerase is proportional to the torsional stress on that polymerase *ϵ* (i.e. ignoring DNA buckling), then Ω = *Nϵ*, and Equation 16 can be written as a function of *N* and *ϵ* as

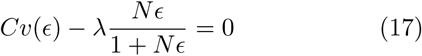

The above equation sets the value of torsional stress *ϵ* for each polymerase and in turn their velocity. Figure 2d shows the contribution of each term in Equation 17 as a function of *ϵ*.

Increasing the polymerase loading rate *R*_*int*_ increases *N*. However, the rate of removal of supercoiling density due to topoisomerase actions saturates with increasing *N* (Figure 2e). The value at which this saturation occurs sets the value of torsional stress on each polymerase and thereby their average velocity. The dashed curve shows the smallest possible *ϵ* that satisfies equation 17 when *N* → ∞ and in turn sets the values of elongation rate for large *R*_*int*_ in Figure 2c. Because of the saturation of the rate of topoisomerase action, the average polymerase velocity can be smaller than the maximum allowed velocity *v*_*max*_. Other choices for the functional form relating the rate of topoisomerase action to Ω(*t*) such as *R*_*topo*_ ∼ *Cnst*. or *R*_*topo*_ ∼ Ω(*t*) will not result in a velocity that saturates to a value other than *v*_*max*_ for large *N* in disagreement with empirical observations where average polymerase elongation velocities both plateau as a function of initiation rate but do so at a value less than *v*_*max*_ [4, 17].

Next, we computed the fluctuations in the number of mRNA molecules. We computed the Fano factor of the number of mRNA molecules (variance divided by mean) over the entire duration of the simulation after steady-state was reached. Surprisingly, the Fano factor deviates from simple Poisson statistics at sufficiently high initiation rate and low rate of topoisomerase action (Figure 2e). This is because as polymerases move along the gene, they interact with each other and change their separation distances (referred to as clustering) from the initial exponentially-distributed separation distances set by the Poisson loading process (Figure 2f). With interactions, the distribution of separation distances peaks at a non-zero value and has a narrower range as evident in Figure 2f. This effect is larger for a lower rate of topoisomerase action because the interactions are mediated by the accumulation of supercoiling density. Clustering of polymerases due to interactions is a plausible explanation for the universally observed ‘bursting’ dynamics of gene expression and the relationship between the average number of transcripts observed in individual cells and the fluctuations of the number of transcripts across cells in a population [9]. The model shows a qualitative relationship between average polymerase velocity and fluctuations in mRNA production (Fano factor, i.e ‘bursting’) consistent with experimental observations. In these experiments, decreasing the rates of topoisomerase action (*λ* in our model) leads to a decrease in the average polymerase velocities and an increase in the levels of mRNA fluctuations (see Figures 3 and 7 in [17]). The same behavior is also displayed by our model (Figure 2c and 2d). Additionally, previous experiments have shown that fluctuations in mRNA production (‘bursting’) is linked to polymerase separation (‘clustering’) [18, 19] by directly observing polymerases as they elongate. Our model is the first theoretical demonstration of this link (Figures 2d and 2f).

**FIG. 3:**
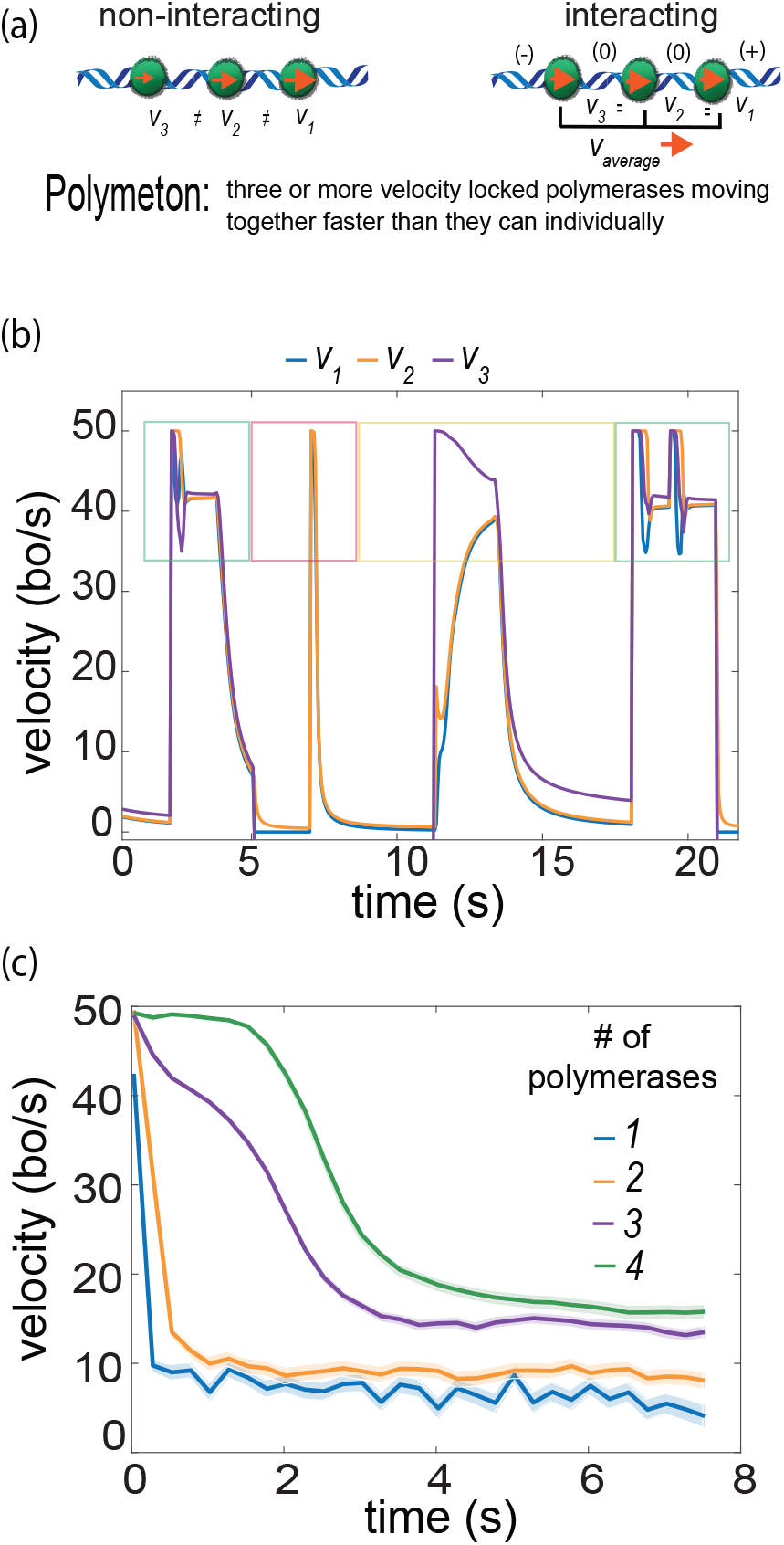
Supercoiling mediated interactions lead to cooperative elongation rates through the creation of Polymetons. (a) Supercoiling density differences across different polymerases lead to velocity differences that in turn change the polymerase separation distances. Polymetons are velocity-locked groups of three or more polymerases where supercoiling density does not accumulate for the middle polymerase. (b)Representative velocity trajectories that show cycles of elongation and stalling due to supercoiling density accumulation and release via topoisomerase action. Three polymerases moving as a polymeton are initially present on the gene labeled with numbers that increase from the termination site to the start site (green square).At approximately t=5s, the leading polymerase reaches the termination site and is removed leaving behind two polymerases that rapidly stall (orange square). Initiation of a new polymerase at the start site increases the elongation rate of the two stalled polymerases mediated through supercoiling-induced interactions (yellow square). Finally, the three polymerases again form a polymeton and elongate with approximately similar velocities (green square on the right). (c) The average elongation rate as a function time following topoisomerase action binned by the number polymerases present on the gene. There is a clear jump when the number of polymerases present changes from two to three showing the formation of polymetons.

To gain an intuition for how clustering occurs, consider the dynamics of two neighboring polymerases. The relative distance between the two polymerases changes as

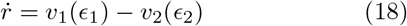

where the velocities of each polymerase is determined by equation 12. The torsional stress experienced by the first and second polymerase is a function of supercoiling density in front of, in between, and behind the two polymerases *σ*_*F*_, *σ*_*M*_, *σ*_*B*_ respectively. In particular, *ϵ*_1_ ∝*σ*_*F*_ − *σ*_*M*_ and *ϵ*_2_ ∝*σ*_*M*_ − *σ*_*B*_. Because supercoiling density always accumulates in the front and all polymerases move in the same direction, *σ*_*F*_ > *σ*_*M*_ > *σ*_*B*_. Importantly, the supercoiling density between the two polymerases is inversely proportional to their separation distance, *σ*_*m*_ ∝ 1*/r*. If the separation distance between two polymerases is large then *σ*_*m*_ is small which in turn generally implies *ϵ*_1_ > *ϵ*_2_ and *v*_1_ < *v*_2_. Therefore, in the case of large separation, 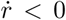 and the separation distance between the two polymerases shrinks. Conversely, if the separation distance between the two polymerases is small then *σ*_*m*_ is large which in turn generally implies *ϵ*_1_ < *ϵ*_2_ and *v*_1_ > *v*_2_ and an increasing separation distance 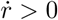. Taken together, as polymerases move along gene, because of the interactions, they converge to a preferred separation distance (Figure 2f) and then move with a constant velocity (Figure 3b). This is the first theoretical description of a natural mechanism for polymerase clustering [18, 19].

Surprisingly in our simulations, we observed that a minimum number of three interacting polymerases moves a larger distance before stalling compared with one or two polymerases, as shown in Figure 3b. To gain an intuition for why a minimum of three interacting polymerases are required for sustained motion, consider how stress accumulates as one polymerase moves along the gene. We can write an equation for the rate of change of the velocity of the polymerase 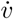 in terms of the dynamics of the local stress by applying a simple chain rule to the stress-dependent velocity (eq.12)

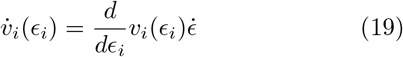

An isolated polymerase moves at velocity *v*_*max*_ initially. As it moves along the gene, the torsional stress across the polymerase accumulates at the rate that is proportional to its velocity, 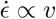. Substituting this into Eq. 19, implies 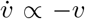. Therefore, the velocity of the polymerase decays exponentially to zero. When topoisomerase relieves the stress, the polymerase can start to move again. When two polymerases move along the gene, their velocities converge to the same value as described above. As with the case of an isolated polymerase, the two polymerases also accumulate torsional stress at a rate proportional to their velocity. This occurs even though supercoiling does not accumulate in the region between the two polymerases because the negative supercoiling density generated by the leading polymerase cancels the positive supercoiling density generated by the trailing polymerase. However, supercoiling density does accumulate outside the two polymerases because there is no cancellation. Therefore, the velocities of the two polymerases also decay exponentially as is the case with a single polymerase (shown in example traces in Figure 3b).

This picture changes qualitatively with the addition of a third polymerase. The velocities of the three polymerases also converge to a shared constant value as they move along the gene. However, the supercoiling density generated by the middle polymerases is exactly canceled by the two polymerases on either side. Therefore, the middle polymerase has no torsional stress accumulation. This puts the middle polymerase in a privileged position. As the outer two polymerases slow down from the accumulation of stress, the middle polymerase continues to move. When this happens, the middle polymerase accumulates torsional stress because there is no exact cancellation of supercoiling density from the other two polymerases. Importantly, the accumulation of supercoiling density in the middle relieves torsional stress on the outer polymerases, resulting in their collective motion. Taken together, three polymerases can sustain their motion for significantly longer periods of time before stalling (as shown in the traces in Figure 3b).

In summary, our results show that there is an emergent phenomenon where three or more polymerases can sustain their collective motions for a longer period of time than one or two polymerases. We will refer to three or more interacting polymerases undergoing sustained motion as polymetons. Polymetons emerge in our system following a topoisomerase action when three or more simultaneously elongating polymerases move at near-constant velocity (Figure.3b). While the precise form of this behavior depends on the details of the model, the qualitative behavior that middle polymerases occupy a privileged position and can assist in sustaining the motion of the group should not depend on the precise form of the model.

### Torque dependant initiation

Next, we incorporate torque-dependent initiation into the model. Up to this point, polymerase initiation in the model occurs stochastically as a Poisson process with a constant rate *R*_*int*_. However, there is experimental evidence that the initiation rate should depend on the supercoiling density at the promoter site. First, there are in vivo measurements showing that increasing the level of negative supercoiling at a promoter by inducing the production of neighboring genes can alter its output [5, 21, 36, 37]. Second, in vitro single-molecule experiments have directly measured promoter unwinding kinetics as a function of the torque in DNA [31] demonstrating that initiation is sensitive to the torque at the promoter.

Polymerase initiation is basically an ordered process of polymerase binding to DNA at the promoter site, unwinding of the DNA, and polymerase leaving the promoter (referred to as clearance). Polymerase-DNA binding follows standard chemical kinetics whereby the binding rate increases proportionally with polymerase concentration and promoter affinity, *K*_*o*_, which captures the intrinsic affinity of polymerase for the promoter, *k*_*b*_ = [*RNAP*]*K*_*o*_. The unbinding rate of polymerase from the promoter, *k*_−*b*_, is not sensitive to polymerase concentration [38]. Precise kinetic observations [31, 38] have measured these rates for specific promoters and could thus be used as inputs into a model of initiation. In our model, we explicitly vary *k*_*b*_ to capture the behavior of genes with different promoter affinities and polymerase concentrations. Inducing or repressing a gene corresponds to varying *k*_*b*_.

Following promoter binding, the polymerase locally unwinds the DNA at the promoter site. This step has a strong dependence on the torsional state of DNA and can become the rate-limiting step in initiation. Precise characterization of promoter unwinding and clearance by polymerases for varying levels of DNA supercoiling has been made [31]. The rate of unwinding depends on the level of torque at the promoter site: the rate of unwinding decreases as torque is increased following a simple Arrhenius law form as measured by [31]. Consequently, we model the unwinding rate *k*_*u*_ as

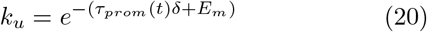

*δ* captures the dependence of the unwinding rate on the torque *τ*. *E*_*m*_ captures the strength of the promoter and can vary from one promoter to another. *E*_*m*_ sets the value at which torque unwinds specific promoters and allows transcription. Some promoters require negative torque to unwind and allow transcription (referred to as weak promoters) whereas strong (consensus) promoters can unwind for positive values of torque [31] (rrnBP1 and lacCONS promoters respectively).

Importantly, the torque at the promoter site *τ*_*prom*_(*t*) is given by the supercoiling density which is in turn set by the polymerases as they move along the gene and interact (Eq. 10). Promoter clearance occurs rapidly following the promoter unwinding [38] and thus is ignored here. Collectively, our model of initiation is composed of a two-step process of reversible polymerase-DNA binding followed by irreversible promoter unwinding which results in initiation (Figure 4a). Here we utilize kinetic unwinding data of the Lac promoter [31] to model a strong promoter requiring no free parameters (see Figure 4b). The inferred values are *E*_*m*_ = −9 and *δ* = 2 (values for *k*_*±b*_ are given in [31]). The values for the weak promoter rrnBP1 are the same but with *E*_*m*_ = 5. This allows us to make a direct comparison between torsion-dependent initiation and gene expression for the same promoter [10] from experiments.

**FIG. 4:**
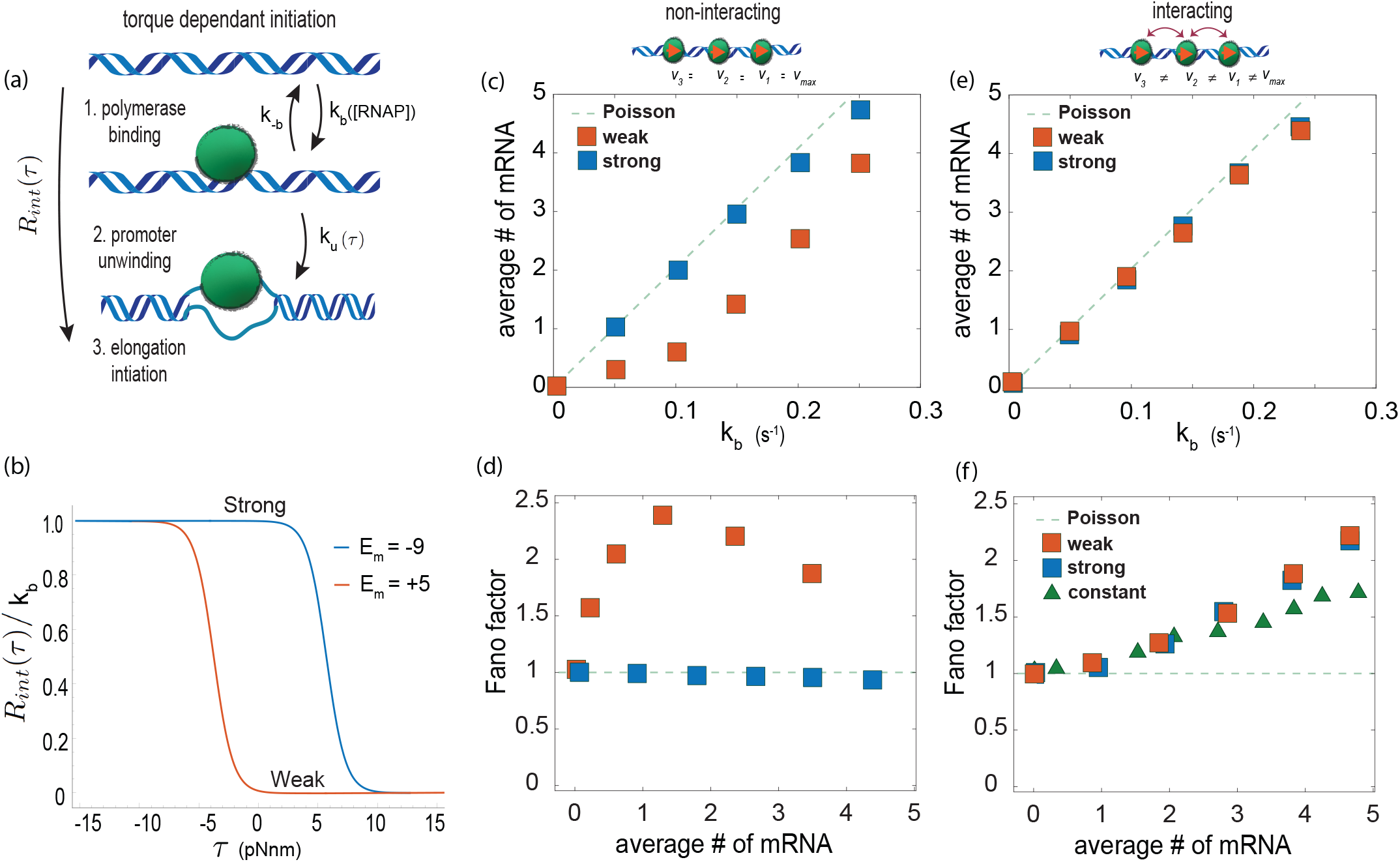
Torque-dependent initiation alongside super-coiling mediated interactions recapitulates the expected mean mRNA production rate and its fluctuations. (a) Torque-dependent initiation is a two-step process: a reversible polymerase binding step that does not depend on torque but on the free RNA polymerase concentration, followed by an irreversible promoter unwinding step which depends on the torque at the promoter site. (b) A weak promoter (*E*_*b*_ > 0) is more sensitive to the torque at the promoter site than a strong promoter (*E*_*b*_ < 0) requiring negative torque to initiate transcription. (c-d) Mean expression and expression fluctuations (Fano factor) for torque-dependent initiation with increasing polymerase binding rate *k*_*b*_ both in the absence of supercoiling mediated interactions between polymerases. The weak promoter exhibits a production rate that scales non-linearly with *k*_*b*_ (c) and non-monotonic fluctuations in expression (d). (e-f) Same as in (c-d) but with the addition of super-coiling mediated interactions between the polymerases. Both strong and weak promoters exhibit linear production rates in *k*_*b*_ (e) with fluctuations that increase monotonically with the average production rate (e). The green triangles in (f) show the fluctuations when interactions are present but the initiation rate is not dependent on torque.

The torque-dependent initiation rate constructed above can be incorporated into our existing framework (Figure 4). We first examine the role of torque-dependent initiation on gene output without polymerase interaction. Figure 4c and 4d show that a strong promoter is not affected by the changes in torque at the promoter site generated by the elongating polymerases even if they are not interacting. In this case, the gene mRNA output increases with the binding rate *k*_*b*_ but exhibits Poisson fluctuations (Fano Factor= 1). A weak promoter (Figures 4e and 4f) displays a non-linear relationship between the average mRNA output and the binding rate *k*_*b*_. However, the fluctuations in the output display a non-monotonic dependence on the mean expression in disagreement with general experimental observations [9] as well as specific observations for the rrnBP1 and lac promoters [10].

Importantly, when the polymerases are allowed to interact as they move along the gene, the dependence of mRNA output and its fluctuation changes. Both weak and strong promoters show a linear dependence of average mRNA output and binding rate *k*_*b*_ (Figure 4e). This output matches what would be expected if polymerase initiation did not depend on the torque but followed a simple Poisson process. The change in the behavior of output is because the torque at the promoter site changes when polymerases interact with each other. In addition, interacting polymerases also change how the mRNA output fluctuates (Figure 4f). Both weak and strong promoters now show super-Poissonian fluctuations with Fano factors that monotonically increase with the average output, again consistent with empirical observations of a super-Poissonian relationship between mean mRNA expression and mRNA fluctuations (see Figure 3 of [10]).

Taken together, these results indicate that torque-dependent initiation alone is insufficient to explain the expected bursting behavior of genes. However, incorporating polymerase (velocity) interactions recovers the expected bursting behavior in genes pointing to fluctuations incurred during elongation as the overriding source of bursting in gene expression. Additionally, the insensitively of strong promoters to negative torque (supercoiling) [31] calls into question the widespread use of initiation as the source of transcriptional bursting used in models [5, 16, 24].

Finally, the interplay between torque-dependent initiation and elongation could be exploited to engineer novel gene regulatory mechanisms or identify existing ones. For instance, imagine two identical genes convergently oriented towards one another both with weak promoters (Figure 5A). The gene that spontaneously initiates transcription first generates positive supercoiling density at the promoter site of the other gene because of the geometry of their arrangement. This supercoiling density in turn changes the initiation rate of the other gene. Therefore, the two genes repress each other resulting in DNA supercoiling-mediated toggle switch.

**FIG. 5:**
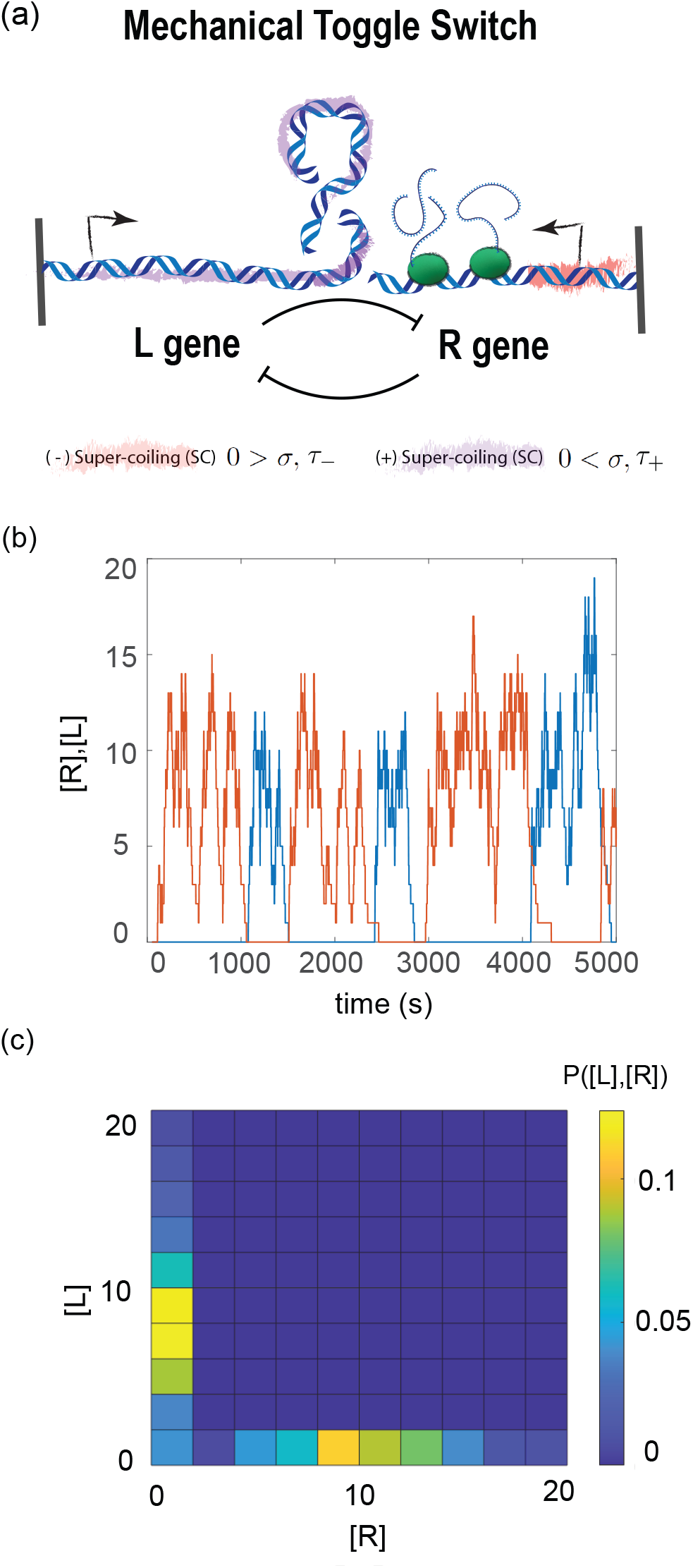
(Mechanical toggle switch demonstrates that DNA supercoiling can be used to engineer interactions across genes without using proteins. a)Two neighboring genes with their promoters oriented towards each other can regulate each other’s expression through supercoiling mediated initiation and elongation resulting in a mechanical toggle switch. (b) Simulated production levels of two identical genes with weak promoters with the orientation shown in (a) as a function of time. The two genes mutually repress one another leading to alternating periods of mRNA production by the [L] and [R] gene. (c) Alternating periods of mRNA production leads to a bi-modal distribution of mRNA expression levels.

To demonstrate this effect, we constructed a system of two convergently oriented, identical genes with torque-dependent initiation and weak promoters. This system demonstrates bi-stable expression where when one gene is on, the other is off (Figure 5b-c). Importantly, the bistability of this system is not set by proteins and therefore the timescale of the oscillations are independent of protein lifetimes. This simple system highlights the potential of novel forms of gene regulation mediated through supercoiling.

## Discussion

The framework introduced here provides a description of the supercoiling density generated during polymerase elongation, the transportation and accumulation of supercoiling density, and its resulting non-local effect on the elongation of polymerases. In addition, we introduced a mechanical model of transcription initiation compatible with our model of elongation. This addition is based on biophysical reasoning and in-vitro observations of torque-dependent polymerase binding and is a crucial element in the construction of a comprehensive framework of supercoiling and transcription. Previous attempts have fallen short of a comprehensive description relying on discrete descriptions of polymerases[15], incomplete models of supercoiling transport due to elongation [16, 24] or models which did not include both torque dependant initiation [15] and elongation [16, 24].

To make the model more computationally tractable some simplifying assumptions were made. First, we assume that supercoiling density diffuses infinitely fast along the gene. This assumption was motivated because the mechanical state of DNA can relax much more quickly compared with the rate at which polymerase moves along DNA. Second, the conversion of DNA twist into writhe (bending) was not explicitly incorporated into our model. Instead, we used a phenomenological formulation of the relationship between supercoiling density and torque which implicitly includes DNA buckling [26]. Similarly, a simple phenomenological framework was used to model the drag on elongating polymerases [23]. Both of these phenomenological models are motivated by physical models of how polymers behave [23, 26]. Third, we constructed a simple model of topoisomerase action. Our model was motivated by empirical observations of topoisomerase kinetics acting on supercoiled DNA plasmids [35]. The phenomenological form used in our model follows exactly the observations made in [35]. Additionally, alternative forms resulted in behaviors that disagreed qualitatively with experimental observations (Figure S2). Finally, the existence of cooperative elongation kinetics and polymetons was found to be robust under various phenomenological parameters used for DNA buckling, nascent mRNA drag and topoisomerase action (Figure S1).

The twist transport equation developed here (eq.9) can be used in future work where DNA writhe is computationally or analytically incorporated allowing for important effects such as histone occupancy dynamics to be included. These effects are particularly important for understanding gene expression dynamics in eukaryotic cells. The current form of the model is most directly applicable to studying the effects of DNA supercoiling on gene expression in prokaryotes.

Many model outcomes have been tested directly or are testable in future experiments. First is the observation that the rate of topoisomerase action determines the average elongation velocity of polymerases. It has been shown previously that the average elongation velocity is below the maximum elongation velocities of polymerases because of topoisomerase action [4, 17]. Additionally, it was previously shown that stopping initiation can slow already elongating polymerases [4]. Our model correctly displays these same behaviors (Figures 2c,2e and S3). The second observation is that the rate of topoisomerase action can control fluctuations in gene expression. Specifically, lowering the rate of topoisomerase action simultaneously lowers the average elongation velocity of polymerases while increasing fluctuations in mRNA production (Figures 3 and 7 in [17]). Our model shows the same behavior (Figures 2c and 2d). The third observation is a super-Poissonian and monotonic relationship between the mean and the fluctuations in the number of mRNAs (see [9] and Figure 3 in [10]). The model correctly predicts these three observations and links them all to the non-local interactions between polymerases mediated by supercoiling. We also predict that the extent of fluctuations in mRNA production is predominantly set by torque-dependent elongation as opposed to initiation (Figure 4). This prediction can be tested by using reporters for directly observing mRNAs as they are being transcribed [18, 39] where one reporter is close to the promoter site while the other reporter is close to the end of the gene. The signals from the two reporters can be used to infer the extent of fluctuations generated by initiation versus elongation. A single reporter system of this kind has been previously used to confirm the relationship between bursting and clustering [18, 19]. Finally, similar reporter systems can be used to directly observe formation polymetons.

In summary, we show that non-local interactions between polymerases emerge from DNA supercoiling and lead to cooperative elongation rates, clustering of polymerases, and gene expression bursting, consistent with empirical observations of cooperative elongation kinetics [4], bursting [9] and clustering [18, 19]. This is the first theoretical effort to link bursting, clustering and elongation rates together in a consistent framework. A collective phenomenon, referred to as Polymetons, emerges whereby groups of three or more polymerases can sustain their elongation due to the privileged status of the interior polymerase. The addition of torque-dependent initiation alone is insufficient to reproduce observed relationships between the mean levels and fluctuations in gene expression. We found that including both torque-dependent initiation and elongation is sufficient to match experimental observations. This result calls into question the validity of using models of torque-dependent initiation alone that do not also incorporate interactions to explain gene expression fluctuations. Finally, we constructed an example where mechanical epigenetics can be used to build novel synthetic regulatory circuits using two neighboring genes to form a mechanical toggle switch.

During the development of this article, two related works with some overlapping results appeared [40, 41]. The first article [40] used an existing model from [15] and the second article makes no explicit connection to DNA mechanics and is largely focused on explaining the recent experimental results that increasing initiation rates can increase elongation rates [4]. In both articles, the connection between elongation kinetics, clustering, and gene expression fluctuations is not made. In fact in [41], it is stated that their model hinders the formation of clustered convoys of polymerases [19]. Importantly, both articles do not report the existence of polymetons and do not examine the effects of torque-dependent initiation on gene expression fluctuations.

In conclusion, we have shown that DNA supercoiling serves as a powerful mediating force between polymerases by altering initiation and elongation kinetics. These effects are inescapable physical attributes of transcription and offer a widespread non-local feedback mechanism between polymerases. This framework challenges the traditional decoupling of transcriptions initiation and elongation and its implications for gene expression fluctuations. Understanding the role that DNA mechanics plays in gene expression can provide new insights into how genes are regulated through mechanical epigenetics.

The authors acknowledge funding from NIH NHLBI R01HL158269 grant.

## SUPPORTING INFORMATION APPENDIX (SI)

### Simulation details and parameters

Simulations were conducted in MATLAB. Euler’s method was used for the numerical integration of the twist angles given in equation 7 of the main text. Integration steps of size Δ*T* = 1*/*200(*s*) where used. The same method was applied for the integration of polymerase motion. See [15] for more details. Stochasticity for the initiation of polymerases, degradation of produced mRNA, and topoisomerase action were modeled as Poisson processes. This was implemented in the simulation by allowing for each process to occur within each time-step with a probability specified by the rate for that process.

### Model predictions are robust to the parameterization of drag

Simulation parameters for stress-dependent polymerase velocity (equation 12 of main text) were *v*_*max*_ = 50, *κ* = 1*/*2*pN*^−1^, *ϵ*_*c*_ = 12*pN*. The drag coefficient associated with polymerase rotation (equation 5 of main text) has a constant term *γ* and a term that increases with the distance *x*_*i*_ of the polymerase from the transcription start site. This form follows from polymer biophysics [23]. However, the precise parameter values for mRNA and polymerase have not been empirically established. *γ* = 10^−1^, *η* = 10^−4^, *α* = 2 were used unless otherwise noted. Figure S1 examines the impact of varying these parameters on the main results of the paper. Figure S1a shows that for any parameter values that result in large drag (*ηx*^*α*^ >> 1) the velocity plateaus at a value less than *v*_*max*_ = 50. This is because a large drag causes significant supercoiling density build-up that impacts the elongation velocities. However if (*ηx*^*α*^ < 1) supercoiling density does not build up and elongation velocities can proceed at rates similar to *v*_*max*_. The size of *ηx*^*alpha*^ (and thus the drag associated with rotation) is dependent on the length of the gene. Consequentially, even if the values for *α* and *η* are small for long enough genes supercoiling density will eventually build up and impact elongation velocities. All values of the parameters *α* and *η* generate a mono-tonic relationship between the average mRNA number and mRNA fluctuations (Fano factor) as shown in S1b. These results show that the model robustly generates the correct behavior for a wide range of parameter values.

### Phenomenological form of the torque of supercoiled DNA

DNA mechanics relax on time-scales much faster than polymerase motion [42]. Due to this fact it is not necessary to explicitly simulate the diffusion of twist between polymerases. Instead we can simply numerically integrate the twist angles of equation 7 of the main text assuming infinitely fast propagation of twist relaxation between polymerases. Thus, throughout the article we employed a steady-state relationship [26] between the level of supercoiling density and corresponding torque in DNA *τ* (*σ*). Our framework allows for buckling to change the torque inside DNA as a function of the supercoiling density. Following the phenomenological approach given by Marko [26] the torque in a given piece of DNA held at a constant force *f* is specified by the supercoiling density as

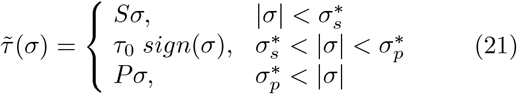

where the coefficients *S, τ*_*o*_, *P* and transition values 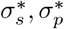 are given by DNA mechanical constants and are a function of applied force (see [26]). The average force *f* inside DNA sets the transition values. Throughout the main text a value of *f* = 1*pNnm* was used. This is a sensible value given the basic polymeric nature of DNA at these scales. To check the robustness of the results of the main text against varying the force *f* we ran the simulation at both *f* = 1*/*10*pN* as well as *f* = 10*pN*. Both values show similar results to simulations run with *f* = 1*pN* (see figure S1).

### Different phenomenological forms of topoisomerase action

As mentioned in the main text, we tried multiple phenomenological forms relating Ω(*t*), the overall supercoiling density in the system (see equation 13 of main text), to the rate of topoisomerase action: a constant rate *λ* independent of Ω(*t*); a rate proportional to Ω(*t*); and a rate that approaches zero with increasing Ω(*t*) (equation 14 of main text). The rate that approaches zero with increasing Ω(*t*) was used throughout the main text. This was because it was the only form that produced results that matched empirical observations for both the elongation velocity and the relationship between average mRNA output and mRNA fluctuations. Topoisomerase action which occurred with a constant rate, as well as an action that occurred with rates linearly proportional to Ω(*t*), were also tried. The results are shown in Figure S2 for the average elongation rates as a function of initiation rates *R*_*int*_ and fluctuations in mRNA production as a function of the average number of mRNAs. Figure S2a shows that a constant rate of topoisomerase action or one that scales linearly with Ω(*t*) both limit the average velocities below *v*_*max*_. However, both of these forms also demonstrated non-monotonic relationships between the rate of initiation *R*_*int*_ and average velocity (Figure S2a) for varying rates *λ*. Additionally, both forms showed a non-monotonic relationship between average mRNA output and mRNA fluctuations (Figure S2b). These results collectively led to the elimination of these phenomenological forms of topoisomerase action for our model.

**FIG. S1:**
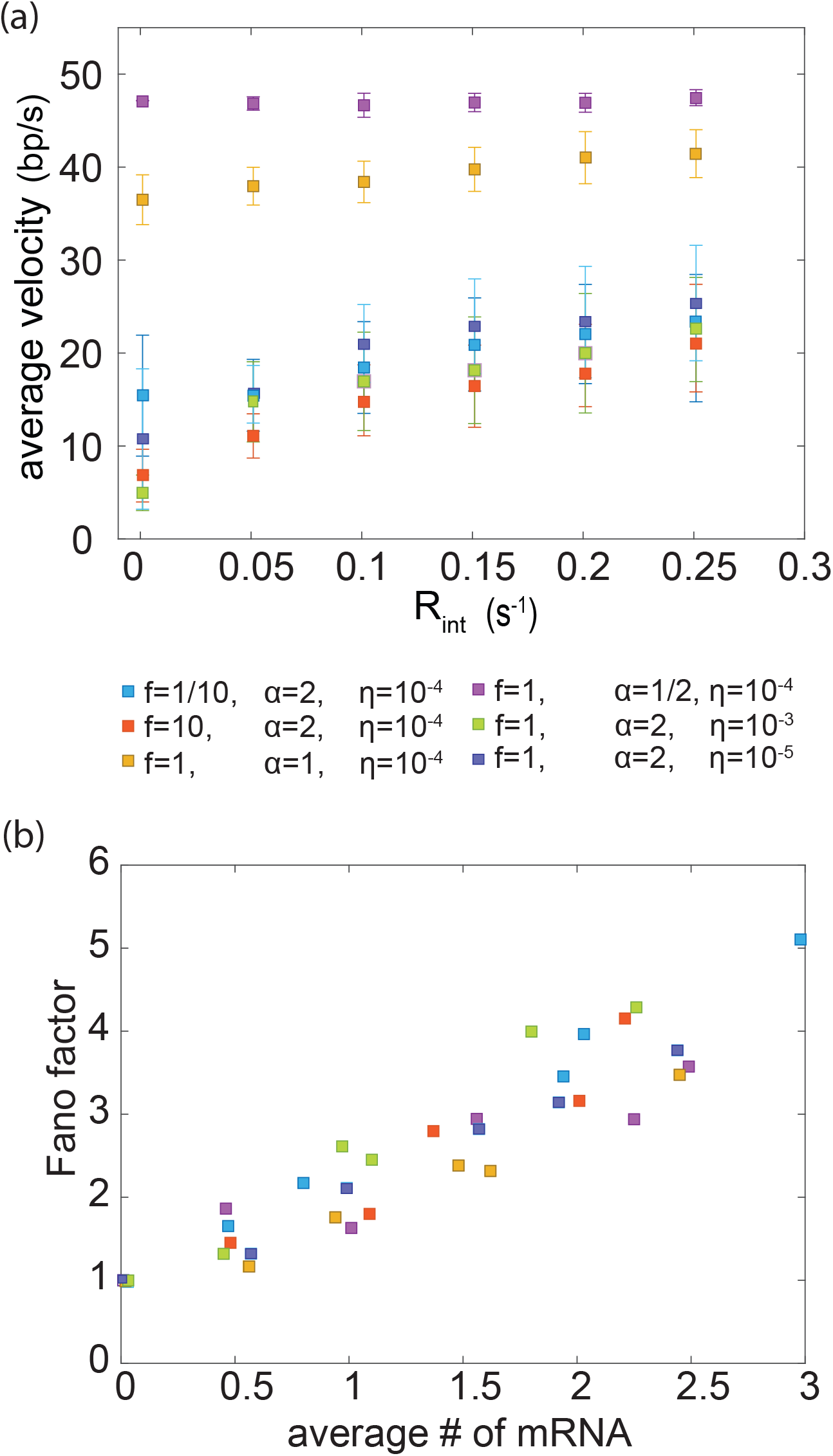
Elongation velocities and the relationship between mean mRNA output and mRNA fluctuations are robust to changes in simulation parameters. (a)The average elongation rate plateaus at a value less than the maximum elongation rate as long as sufficient supercoiling exists at any value for DNA force *f*. If the drag associated with polymerase rotation is not significant (*η, α* are small) then only long genes will create significant supercoiling build up. (b) All parameters values correctly demonstrate a monotonic relationship between the mean mRNA output and mRNA output fluctuations.

**FIG. S2:**
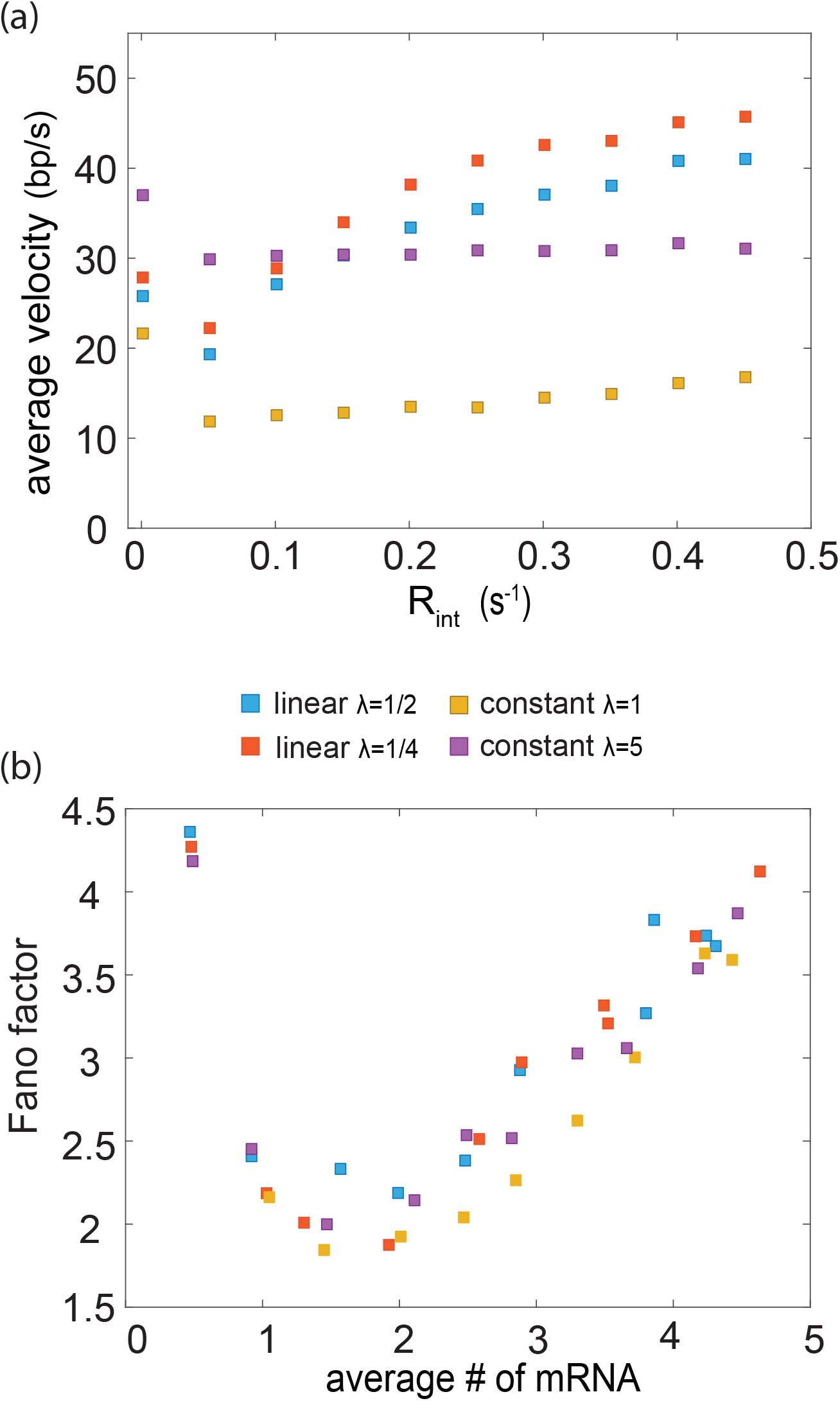
Constant rates of topoisomerase action or rates that linearly depend on supercoiling density do not recapitulate empirical observations. (a)The average elongation rate shows a non-monotonic dependence on the initiation rate for both a constant and linear rate of topoisomerase action. (b) Both a constant and linear rate of topoisomerase action result in a non-monotonic relationship between the average mRNA output and the fluctuations in mRNA. These two results demonstrate that neither a constant nor a rate of topoisomerase action that linearly depends on supercoiling density can recapitulate empirical observations and were therefore not used in the model.

**FIG. S3:**
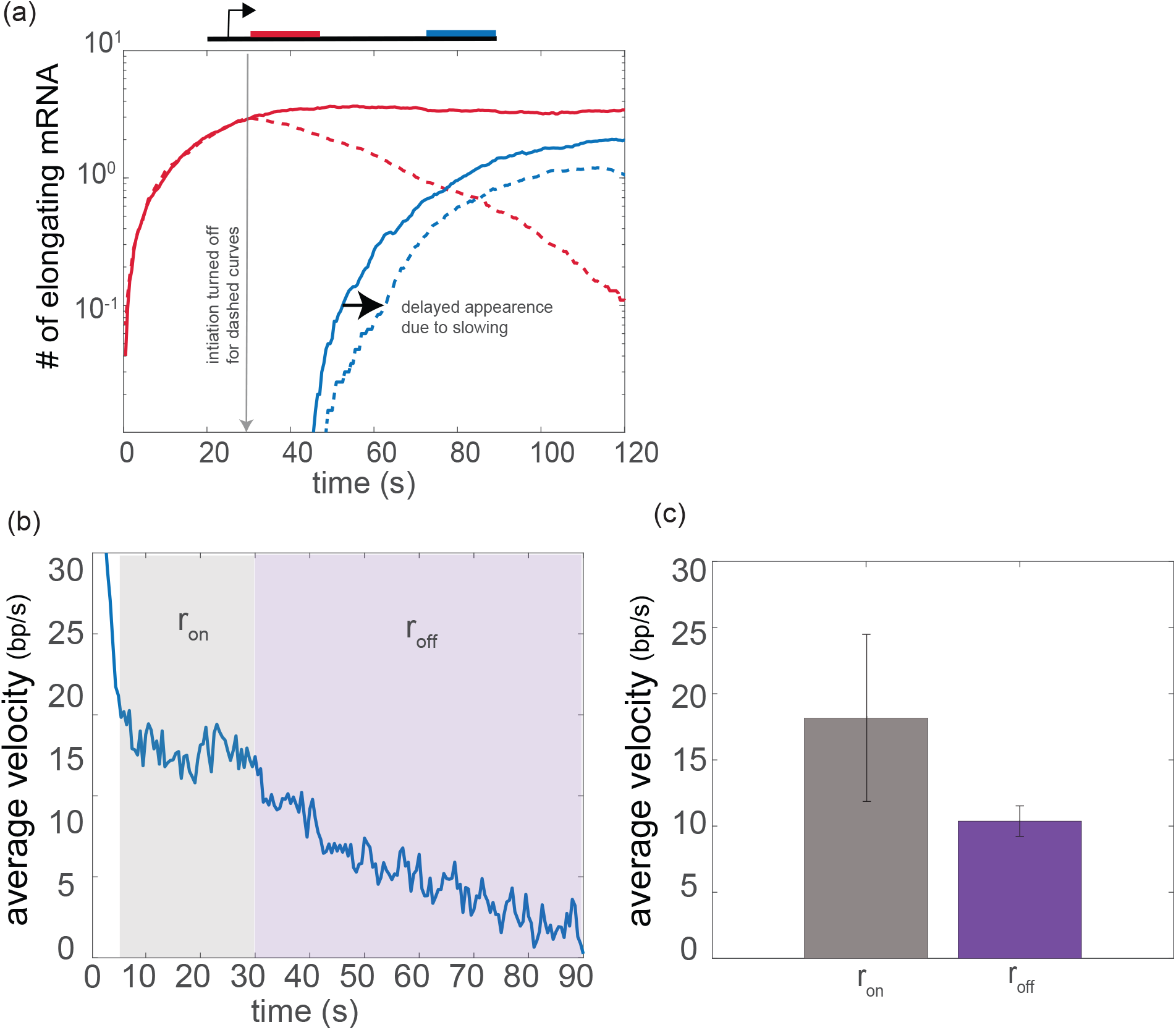
Stopping polymerase initiation leads to the slowing of already elongating polymerases in agreement with experimental observations reported in [4]. (a) We simulated polymerase elongation for a gene with one reporter close to the promoter site (red) and another reporter located further downstream (blue). After initiation, elongating polymerases first pass through the red reporter than the blue reporter. We can determine the average elongation velocity of polymerases between the two reporters by measuring the time that elapses from the onset of the signal from the red reporter to that of the blue reporter (*velocity* = *distance/time*). Two experiments are simulated. The first has initiation turned on at *t* = 0 and left on (solid line) and the second has initiation turned on at *t* = 0 and then turned off at *t* = 30*s* (dashed line). The shift in the dashed blue curve from the solid blue curve indicates that turning off initiation causes already elongating polymerases to slow. (b) The average velocity of all elongating polymerases while initiation is kept on *r*_*on*_ followed by the average velocity of all elongating polymerases after initiation is turned off *r*_*off*_ as a function of time. (c) The average velocity of elongating polymerases during the periods of *r*_*on*_ and *r*_*off*_. Elongating polymerases require the initiation of additional polymerases to maintain their velocity.

